# Demographic impacts of low- and high-intensity fire in a riparian savanna bird: implications for ecological fire management

**DOI:** 10.1101/2022.12.10.519856

**Authors:** Niki Teunissen, Hamish McAlpine, Skye F. Cameron, Brett P. Murphy, Anne Peters

## Abstract

1. Climate change is driving changes in fire frequency and intensity, making it more urgent for conservation managers to understand how species and ecosystems respond to fire. In tropical monsoonal savannas – Earth’s most fire-prone landscapes – ecological fire management aims to prevent intense wildfires late in the dry season through prescribed low-intensity fire early in the dry season. Riparian habitats embedded within tropical savannas represent critical refuges for biodiversity, yet are particularly sensitive to fire. Better understanding of the impact of fire – including prescribed burns – on riparian habitats is therefore key, but requires long-term detailed post-fire monitoring of species’ demographic rates, as effects may persist and/or be delayed.
2. Here, we quantify the multi-year impacts of prescribed low-intensity and high-intensity fire on the density, survival, reproduction and dispersal of the threatened western purple-crowned fairy-wren (*Malurus coronatus coronatus*), in an exceptionally well-studied individually-marked population.
3. Following low-intensity fire, bird density was reduced in the burnt compared to adjacent unburnt riparian habitat for at least 2.5 years. This was a result of reduced breeding success and recruitment for two years immediately following the fire, rather than mortality of adults or dispersal away from burnt habitat.
4. In contrast, a high-intensity fire (in a year with low rainfall) resulted in a sharp decline in population density 2-8 months after the fire, with no signs of recovery after 2.5 years. The decline in density was due to post-fire adult mortality, rather than dispersal. Breeding success of the (few) remaining individuals was low but not detectably lower than in unburnt areas, likely because breeding success was poor overall due to prevailing dry conditions.
5. Hence, even if there is no or very low mortality during fire, and no movement of birds away from burnt areas post-fire, both low- and high-intensity fire in the riparian zone result in reduced population density. However, the mechanism by which this occurs, and recovery time, differs with fire intensity. To minimise the impacts of fire on riparian zones in tropical savannas, we suggest employing low-intensity prescribed burns shortly after the breeding season in years with good rainfall.

## INTRODUCTION

Fire is an important disturbance and selective pressure, shaping ecosystems worldwide (Nimmo et al. 2021, Gonzalez et al. 2022). However, the frequency, intensity, and size of fires is increasing around the globe, due to anthropogenic land use and climate change causing increasingly warm and dry conditions (Bowman et al. 2009, Wu et al. 2021). A single fire can dramatically alter key resources and the abundance and diversity of species for decades to follow (Gonzalez et al. 2022). As a result, changing fire regimes are transforming terrestrial ecosystems and posing a threat to biodiversity (Kelly et al. 2020, Gonzalez et al. 2022). This is occurring through increased stochastic and sporadic extreme events such as mega-fires (Nimmo et al. 2021), but also in the landscapes traditionally shaped by annual fire seasons, such as monsoonal savannas. These are characterised by a wet season, associated with a build-up of fuel, which cures over the following dry season, promoting high fire frequencies (Andersen 2021). To reduce the risk of frequent and large, high-intensity wildfires, ecological fire management is used extensively in such areas (Russell-Smith et al. 2013, Andersen 2021). This typically involves prescribed burning early in the dry season, when fire weather conditions are relatively mild and fires tend to be small, patchy, and of low intensity. Creating a mosaic of patches varying in fire history promotes plant and animal diversity and reduces the extent and frequency of large, high-intensity wildfires late in the dry season (Parr and Andersen 2006, Russell-Smith et al. 2013, Andersen 2021). However, this approach to fire management currently does not sufficiently consider the effect of fire on animal movement within or among habitat patches, or the implications for long-term population persistence (Sitters and Di Stefano 2020).

Riparian ecosystems embedded within fire-prone tropical savanna landscapes are among the most vulnerable in the face of climate change (Tockner and Stanford 2002, Capon et al. 2013). Yet they are also vitally important: they harbour high biodiversity, act as movement corridors, and provide key refugia from climate warming and drought (Woinarski et al. 2000, Capon et al. 2013, Fremier et al. 2015, Krosby et al. 2018). Although riparian zones can also provide a refuge from and buffer against fire, under certain conditions, riparian zones become corridors through which fire can spread (Pettit and Naiman 2007). Ecological consequences of fire in riparian zones are more severe compared to surrounding savanna (Pettit and Naiman 2007, Douglas et al. 2015, Flores et al. 2021). It is therefore of vital importance to quantify effects of fire on riparian zones, and their recovery, to inform management of these key refugia (Pettit and Naiman 2007, Douglas et al. 2015).

Particularly, we need to enhance our understanding of the underlying processes through which animal populations are affected by fire. Direct mortality rates during fire seem to be generally low (^~^3%) (Nimmo et al. 2021, Jolly et al. 2022). However, fire also affects populations indirectly, through habitat change, reducing availability of food and shelter and increasing predation (McGregor et al. 2015, Andersen 2021, Jolly et al. 2022). This may affect survival and/or breeding success (Murphy et al. 2010), resulting in population decline. Animals may also modify their behaviour in response to fire, for example moving to unburnt habitat during or immediately after the fire (Murphy et al. 2010, Pausas and Parr 2018, Nimmo et al. 2019, Nimmo et al. 2021). Hence, apparent population declines may also result from such behavioural adaptations. To identify the mechanisms of fire impacts on populations, it is therefore critical to quantify mortality, movements and breeding success of individuals pre- and post-fire compared to unburnt controls (Driscoll et al. 2010, Nimmo et al. 2019).

Here, we investigate the demographic impacts of low- and high-intensity fire in riparian habitat in the tropical savanna of northern Australia, an important refuge for many species, including birds (Woinarski et al. 2000). Our study species is the western purple-crowned fairy-wren (*Malurus coronatus coronatus*), widely considered a biological indicator of riparian health in the region (Skroblin and Legge 2012). It is a riparian endemic listed as Endangered under Australian national legislation due to extensive habitat degradation by feral herbivores and intense fires (Skroblin and Legge 2012). Our study population has been studied since 2005, with detailed records of mortality, movement and reproduction for all individuals. Utilising a before-after control-impact (BACI) design, we quantify fairy-wren density for 2.5 years before and 2.5 years after a low- and a high-intensity fire, in both burnt and adjacent unburnt riparian habitat. Specifically, we investigate: whether fire caused mortality via direct or indirect effects; the role of movement by birds out of burnt habitat; and the importance of reduced breeding success. Finally, we consider how our findings can inform fire management strategies.

## METHODS

### Study system

Research took place at Annie Creek in Australian Wildlife Conservancy’s Mornington Wildlife Sanctuary (17°31’S 126°6’E) in the Kimberley region of north-western Australia. Here, we have studied a population of the western purple-crowned fairy-wren since 2005. Throughout its range, this fairy-wren is absent from degraded riparian vegetation (Skroblin and Legge 2012). This is largely because of its strong positive association with the riparian tree *Pandanus aquaticus* (henceforth referred to simply as *Pandanus*) which declines in abundance as sites become degraded (Skroblin and Legge 2012).

Purple-crowned fairy-wrens live in groups (a dominant breeding pair plus a variable number of subordinates; Kingma et al. 2010) that occupy territories aligned linearly along waterways and defended year-round (Kingma et al. 2010). Birds can disperse to settle as a breeder or subordinate elsewhere (Hidalgo Aranzamendi et al. 2016). Breeding mostly takes place during the wet season (December-April) (Kingma et al. 2010). Purple-crowned fairy-wrens usually lay 3 eggs (range 1-5), incubation lasts for 14 days, and the nestling period for 13 days (Teunissen et al. 2020). Nest failure is common (78% of nests), mostly due to predation (57% of nests) (Teunissen et al. 2020). Individuals can re-nest multiple times in a season.

### Fire history

Northern Australia is characterised by a monsoonal wet season (ca. December – April), followed by a dry season (ca. May – November), with high fire frequencies late dry season (Andersen 2021). At our study site, fire regimes have been managed by AWC since 2004 for biodiversity conservation using prescribed burning early in the dry season (prior to July) to limit large-scale high-intensity wildfires late in the dry season. This approach to fire management is common throughout northern Australia and follows on from traditional fire management imposed on the landscape by Aboriginal people for millennia (Russell-Smith et al. 2013, Andersen 2021). As a result of prescribed burning at Mornington Wildlife Sanctuary, intense fires burning *Pandanus* in riparian zones are generally rare. However, two recent prescribed fires burnt sections of Annie Creek. Prior to these, fire had not burnt the riparian vegetation along Annie Creek for at least 11 years.

The first fire occurred in February 2015. The preceding 12-month period had rainfall slightly below average (748 mm vs. annual average of 868 mm for 2005-2022; Bureau of Meteorology weather station 2076). The fire was lit at Annie Creek, burnt a relatively small area, and fire weather conditions at the time of the fire were mild, with the wind pushing the fire away from Annie Creek. As a result, the fire was of low intensity, patchy – as is common for fires this time of year – and left small stands of *Pandanus* unburnt. The fire burnt 0.78 km of our study site’s 4.2 km stretch of Annie Creek, corresponding to eight burnt purple-crowned fairy-wren territories, while 35 territories remained unburnt (Fig 1a).

**Fig. 1.**
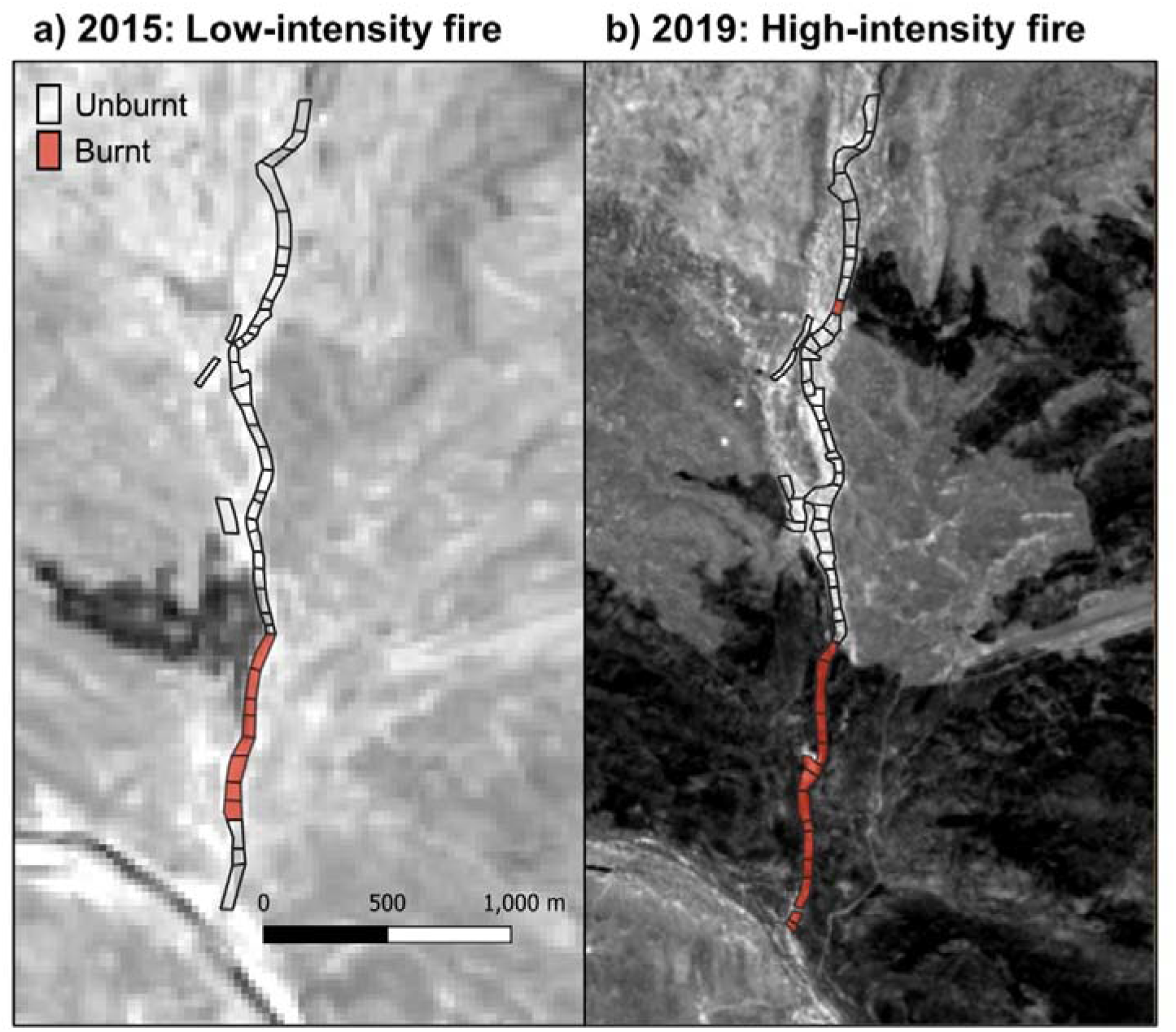
Satellite images showing burnt areas (in black) along Annie Creek indicating the extent and intensity of (a) the small, low-intensity fire in 2015, and (b) the extensive, high-intensity fire in 2019. Polygons depict purple-crowned fairy-wren territories. Polygons highlighted in red were confirmed burnt by ground-based surveys (note that the southernmost burnt territories in (a) had fire trickle through the understory of the riparian strip only, therefore not showing up as clearly as a black scar). One territory further north along the creek was burnt in 2019 (b), but only partly and at lower intensity, and therefore excluded from analyses. Satellite images were obtained from Sentinel Hub, from the (a) Landsat 8 satellite (captured on 18 February 2015), and (b) Sentinel-2 satellite (captured on 19 March 2019).

The second fire occurred in March 2019. The preceding 12-month period had extremely low rainfall (396 mm, the driest 12-month period since records began in 2005). Additionally, the fire was lit >5.5 km away from Annie Creek, but burnt a large area, and travelled to Annie Creek pushed by a strong wind, reaching Annie Creek as a large headfire. As a result of these conditions, despite the time of year, this fire acted more like a late dry season high-intensity fire. Most vegetation within the fire scar was burnt, and small patches of *Pandanus* initially left unburnt in the fire scar subsequently died (Fig. S1). The fire burnt 1.26 km of our study site’s 4.2 km stretch of Annie Creek, including the stretch burnt in 2015, encompassing 13 fairy-wren territories, leaving 36 territories unburnt (Fig 1b). In addition, a short 60 m stretch of creek within a single fairy-wren territory was burnt further north on the creek (Fig. 1b), but at low intensity and on one bank only, making it not directly comparable to the other burnt territories, therefore this single territory was excluded from analyses.

For both fires, the adjacent sections of unburnt creekline (see Fig 1) provide a biologically relevant control, allowing us to assess the effect of fire while controlling for any confounding variables (e.g. variation in climate between years) which impact survival, movement, and breeding success.

### Purple-crowned fairy-wren demography

We monitored density, survival, dispersal, and breeding success of purple-crowned fairy-wrens from 2011 to 2021. All birds along 15 km of Annie Creek and the adjoining Adcock River were uniquely colour-banded and followed throughout life. Each year, we conducted a full census of the population just before the wet season (mid-October to late November; hereafter termed season ‘November’), and just after the wet season (late April to mid-June; hereafter season ‘May’). Additionally, the population was monitored throughout the main breeding season from January to April in 2016-2021 (except 2019). Each season, we recorded the presence, location and social status of all individuals. Density (individuals/km) was calculated as the total number of individuals inhabiting the burnt and unburnt sections of creek (Fig. 1) divided by their length (in km), at the end of each May and November season.

For all individuals present before each fire, we estimated duration of survival and timing of any dispersal. Each individual was considered alive until the last day of the last season in which it was seen in the study population (detection rate per season = 98%; Roast et al. 2020). Dispersal was determined from movements of individuals within the population and, in much rarer cases, emigration. To identify emigrants, we surveyed waterways connected to our study site for banded individuals, using playback of conspecific song recordings (>90% detection probability; Hidalgo Aranzamendi et al. 2016). We surveyed all suitable habitat (i.e. containing *Pandanus*) within 20 km of the study area, and most suitable habitat in a wider 60 km radius (for full details, see Roast et al. 2020). Hence, we can distinguish emigration from death, resulting in highly reliable estimates of survival.

We recorded breeding success over the course of an Austral year (i.e. 1 July – 30 June; to include the breeding peak, December-March). This was expressed as the total number of fledglings, free-flying juveniles recognisable from behaviour, morphological and plumage characteristics, detected in each territory.

### Statistical analyses

All statistical analyses were performed in R version 4.2.1 (R Core Development Team 2022), using the packages ‘lme4’ and ‘lmerTest’.

To test if low- or high-intensity fire affected population density, we used fairy-wren density (individuals/km) for five seasons immediately prior to, and five seasons following each fire. We compared density post- vs pre-fire in the burnt vs unburnt (control) section of creek by running for each fire a linear model with density as response variable, and as independent variables: time period (pre-fire, post-fire), fire treatment (burnt, unburnt), and their interaction.

To analyse whether low- or high-intensity fire affected adult survival, we ran a Cox proportional hazards model, using the ‘coxph’ function from the package ‘survival’. We analysed survival from immediately before the fire (the November field season prior to the fire) up to >2.5 years following fire (November 2017 for the 2015 fire, November 2021 for the 2019 fire). Individuals that were still alive by the end of this period were censored. All individuals present on our Annie Creek study site before the fire were included, except for one bird that was unbanded and could not be individually recognised to monitor survival (low-intensity fire: N = 31 in burnt area, N = 125 in unburnt area; high-intensity fire: N = 51 in burnt area, N = 123 in unburnt area). We ran a separate Cox model for each fire to test whether survival differed between fire treatments (burnt, unburnt), while controlling for the effect of social status (subordinate, dominant) on survival.

To analyse whether low- or high-intensity fire affected dispersal, we used the same approach, sample size, and time period as above for analysing survival. Birds still in their home territory by the end of the period were censored, as were birds that died during the period analysed (as they could not disperse past the moment of death). Again, we ran a Cox proportional hazards model for each fire to test whether the duration birds stayed home for differed between burnt and unburnt regions. We also controlled for bird status, as subordinates are more likely to disperse (Hidalgo Aranzamendi et al. 2016).

Lastly, we analysed how low- or high-intensity fire affected breeding success (range = 0-5 fledglings per territory per year). We compared breeding success over three Austral years before and after each fire. Both fires took place during the period when most breeding takes place. Since the fires likely disturbed that year’s breeding attempts, we considered the wet seasons in which the fires occurred post-fire years. We were unable to include some territories for some years, due to territories splitting, merging, or ceasing to be inhabited. For the low-intensity fire, we had 18 measures of breeding success for burnt territories pre-fire and 24 post-fire, and 88 unburnt territories pre-fire and 101 post-fire. For the high-intensity fire, our sample size included 32 measures for burnt territories pre-fire and 24 post-fire, and 105 unburnt territories pre-fire and 91 post-fire. We compared pre-fire and post-fire breeding success in the burnt and unburnt stretch of creek using generalised linear mixed models with poisson distribution. We included as response variable the number of fledglings produced on a territory in a given year, and as independent variables time period, fire treatment, and their interaction. We included territory ID as random effect to account for repeated measures for territories. Because we found a significant interaction between time period and fire treatment for the low-intensity fire, indicating reproductive success was lower in the burnt area post-fire (see Results), we sought to test how long this effect lasted for. Since no territories in the burnt area produced any fledglings in 2014/2015 (i.e. complete separation of the data; Fig. 4a), we ran a Bayesian GLMM using the ‘rstanarm’ and ‘rstan’ packages. We included the same variables as before except we replaced time period with year. We set priors to a normal distribution with mean = 0 and variance = 10, and variance = 100 for the intercept. We ran three chains of 15,000 iterations, with a thinning interval of 20 and warmup period of 5,000. We visually inspected trace, density, autocorrelation, and posterior predictive plots to confirm convergence of the model. We present posterior mean and 95% credible intervals (CI).

## RESULTS

### Density

After the low-intensity fire, fairy-wren density was reduced in the burnt area, but not in the unburnt control area compared to before the fire (interaction time period and fire treatment; *B* ± SE = −12.04 ± 5.19, t_16_ = −2.32, *P* = 0.03; Fig. 2a; Table S1). This difference reflects a slight decline in density in the burnt area at the time of the fire, and a concurrent increase in density in the unburnt area, while trajectories in both sections of creek appeared similar over the following two years (Fig. 2a).

**Fig. 2.**
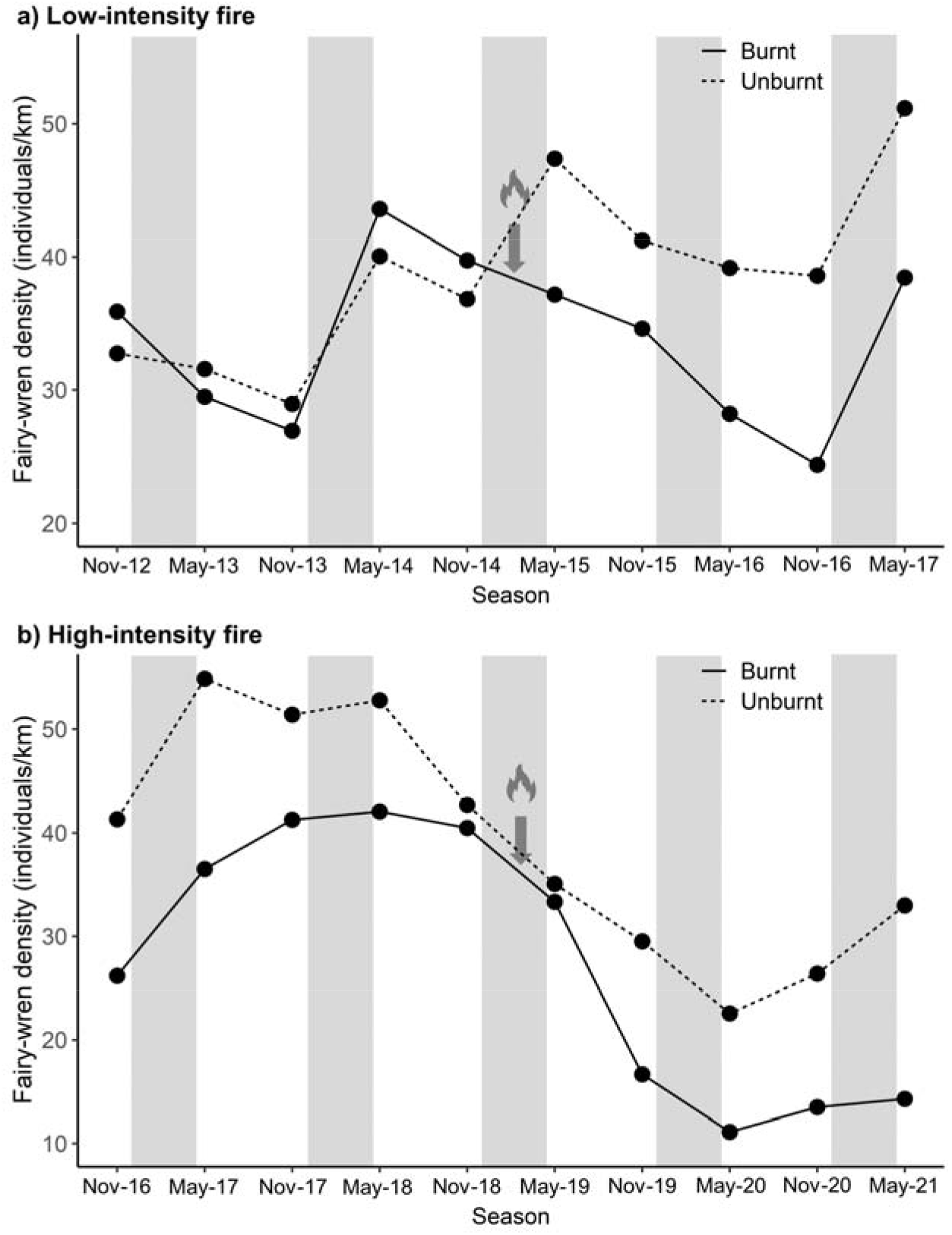
Population density (individuals/km) in burnt (solid line) and unburnt (dashed line) sections of the creek following (a) a low-intensity fire, and (b) a high-intensity fire. Arrows indicate timing of fire. Grey shading indicates timing of wet seasons.

For the high-intensity fire, fairy-wren density was lower post-fire compared to pre-fire in general, and in the burnt compared to the unburnt area (Table S1). However, fairy-wren density did not decrease more in the burnt area compared to the unburnt area (no interaction time period and fire treatment; *B* ± SE = −0.22 ± 6.10, t_16_ = −0.04, *P* = 0.97; Fig. 2b; Table S1). Instead, fairy-wren density in the burnt area seemed to decrease in the six months following the fire (which this statistical model of density cannot formally test for, but which is addressed by the individual survival analyses, see below).

### Survival

Individual survival did not differ between the burnt and unburnt control area following a low-intensity fire (*B* ± SE = −0.29 ± 0.27, z = −1.08, *P* = 0.28; Fig. 3a; Table S2). The high-intensity fire on the other hand significantly reduced survival in the burnt compared to the unburnt area (*B* ± SE = – 0.54 ± 0.58, z = −2.75, *P* < 0.01; Table S2). Survival seemed unaffected shortly following the high-intensity fire, but was clearly reduced in the longer term (2-14 months post-fire) (Fig. 3b).

**Fig. 3.**
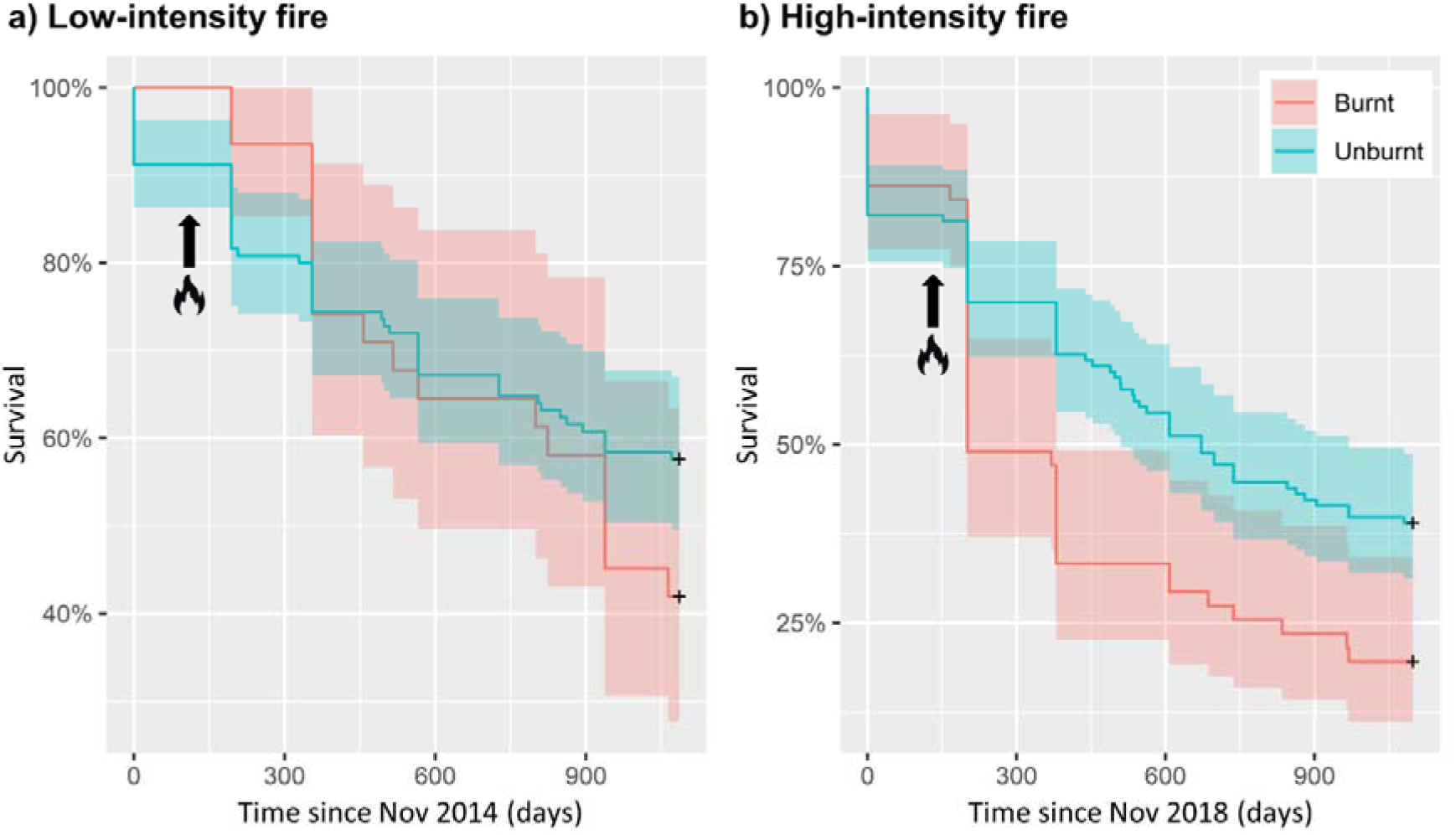
Purple-crowned fairy-wren adult survival was (a) unaffected by a low-intensity fire, but (b) reduced following a high-intensity fire, with survival curves for burnt (red) and unburnt areas (blue) diverging 2 months after fire. Arrows indicate timing of fire. Survival is represented in Kaplan Meier plots. Shaded bands represent 95% CI.

### Dispersal

Individuals from the burnt area did not disperse more often than individuals in the unburnt area following the low-intensity fire (*B* ± SE = −0.40 ± 0.67, z = −1.12, *P* = 0.26; Table S3), or the high-intensity fire (*B* ± SE = 0.22 ± 0.35, z = 0.64, *P* = 0.52; Table S3).

### Breeding success

Just before the low-intensity fire, breeding success was comparable between the burnt and unburnt areas. However, following the fire, breeding success was lower in the burnt compared to the unburnt area (interaction time period and fire treatment; *B* ± SE = −0.94 ± 0.35, z = −2.69, *P* < 0.01; Fig. 4a; Table S4). Breeding success was reduced for two years: in 2014/2015 (interaction year and fire treatment: posterior mean = 7.6, 95% CI = 2.2 – 16.7) and 2015/2016 (posterior mean = 2.6, 95% CI = 0.5 – 5.5). Before and after the high-intensity fire on the other hand, breeding success was comparable between burnt and unburnt territories (no interaction time period and fire treatment; *B* ± SE = 0.10 ± 0.37, z = 0.29, *P* = 0.77; Fig. 4b; Table S4).

**Fig. 4.**
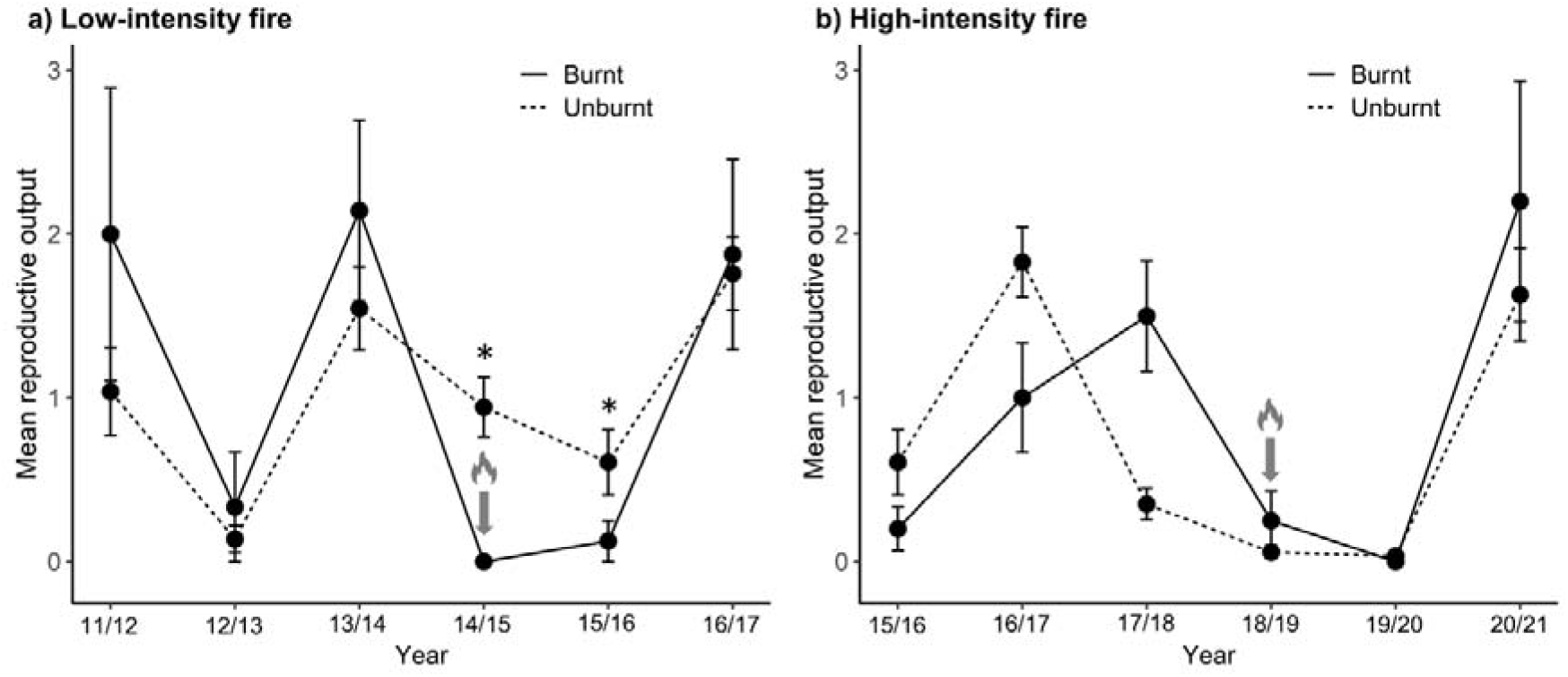
Mean reproductive output (number of juveniles produced per territory) in burnt (solid line) and unburnt (dashed line) sections of the creek following (a) a low-intensity fire, and (b) a high-intensity fire. Error bars represent standard errors. Arrows indicate timing of fire.

## DISCUSSION

We used a before–after control–impact study design to examine population responses of a threatened riparian bird to low- and high-intensity fire. Close monitoring of individually marked birds with complete life-histories provided detailed information on demographic rates, allowing us to identify the mechanisms driving changes in population density. We show that low-intensity fire reduced density in the first months immediately following fire, due to reduced recruitment. In contrast, high-intensity fire had its most pronounced negative effect on density 2-8 months after the fire, due to delayed mortality of adult birds. Since purple-crowned fairy-wrens are a biological indicator for riparian health, our findings can be used to fine-tune fire management practices in riparian habitats in tropical savannas.

Our findings support recent reviews showing that fire causes limited direct animal mortality, and instead impacts survival mostly through indirect effects caused by fire-induced changes to habitat and resource availability (Andersen 2021, Jolly et al. 2022). Our study is unusually robust because we can estimate true mortality with a high degree of accuracy, because we can distinguish death from emigration. Therefore, we can confidently conclude that the low-intensity fire did not result in mortality, immediate or delayed, whereas survival was reduced following high-intensity fire. This reduction in survival became evident >2 months after the high-intensity fire, which degraded habitat substantially (Fig. S1). These habitat changes may increase mortality by reducing resource availability (Valentine et al. 2012), or increasing hunting success of predators because prey are more exposed (McGregor et al. 2015, Hradsky et al. 2017). Additionally, we propose that fire-induced habitat changes likely expose birds to more extreme temperatures due to reduced cover. Riparian areas are characterised by a cool microclimate and provide refugia from climate warming (Krosby et al. 2018). Exposure to increased temperatures can affect animals’ physiology and survival (Sharpe et al. 2019, Conradie et al. 2020), hence increased temperatures after fire may be an important mechanism through which fire indirectly causes mortality. Potential future research will focus on how fire affects microclimate at our study site, and consequences for fitness of fairy-wrens, to reveal additional mechanisms underlying post-fire population dynamics.

The capacity to move away from an active fire or to disperse to unburnt habitat following fire can be important adaptive strategies for avoiding fire impacts (Nimmo et al. 2019, Nimmo et al. 2021). However, purple-crowned fairy-wrens did not disperse in response to fire, nor did breeding groups move their territory to nearby unburnt habitat. This possibly reflects that there was little suitable habitat available for fairy-wrens to disperse to, being restricted to a narrow band of suitable habitat along the creekline (Fig. 1). The fire response of the purple-crowned fairy-wren contrasts markedly with that of its sympatric congener, the red-backed fairy-wren, *Malurus melanocephalus*, which occurs in fire-prone savannas, rather than the riparian zone. Red-backed fairy-wrens dispersed and recentred their territories around unburnt habitat following low- and high-intensity fires (Murphy et al. 2010, Sommer et al. 2018). This response is thought to explain the apparent lack of post-fire mortality (but note that in these studies, individuals were only followed for 1-2 months post-fire). The contrasting fire response of the purple-crowned fairy-wren is consistent with the general notion that riparian species are more strongly affected by fire than savanna species (Pettit and Naiman 2007, Douglas et al. 2015, Flores et al. 2021).

Breeding success of purple-crowned fairy-wrens was reduced immediately after the low-intensity fire, and – to a lesser extent – the following year (Fig. 4a). Reproductive output may have been impacted via several mechanisms. Firstly, since the fire occurred during the main breeding season, active nests would have burnt in the fire (indeed, we observed presumed burnt fairy-wren nests following the high-intensity fire). Secondly, habitat changes after fire likely reduced nest survival by enabling predators to detect nests more easily (e.g. Pittman and Krementz 2016), and/or because parents were more conspicuous when visiting the nest (Martin et al. 2000). Lastly, birds may have been less likely to re-nest, resulting in a shorter breeding season (as in red-backed fairy-wrens; Murphy et al. 2010). Surprisingly, breeding success was not significantly reduced by the high-intensity fire. This is probably because the fire coincided with, and was followed by, poor breeding conditions. Rainfall was unusually low the year the fire occurred (447 mm compared to annual average of 868 mm), and the following year (553 mm). Given that breeding is triggered by rainfall in this species (Aranzamendi et al. 2019), breeding success – including in unburnt habitat – was low for these years (near zero; Fig. 4b), and this probably limited our ability to detect any further depression of reproduction.

Assessing the direct and indirect effects of fire on riparian zones, and their recovery post-fire, is critical to refining fire management practices to protect these key ecosystems (Pettit and Naiman 2007). Our results show that population recovery after riparian fire takes a long time. Following low-intensity fire, reproductive output of purple-crowned fairy-wrens was severely reduced for two breeding seasons. Following high-intensity fire, adult survival was severely reduced for at least a year. As a result, density remained substantially reduced for over two years at least following any fire event (Fig. 2). Since many species rely on riparian vegetation for foraging, dispersal, and to seek refuge from fire and hot temperatures (Woinarski et al. 2000, Capon et al. 2013, Fremier et al. 2015, Krosby et al. 2018), fire in riparian zones is expected to similarly affect a range of species. Recovery of populations after fire generally requires in situ survival as well as immigration from unburnt areas (Shaw et al. 2021, Hale et al. 2022). Intact riparian habitat is however severely fragmented across northern Australia, limiting connectivity between populations (Skroblin et al. 2014), and likely hindering post-fire recovery of many species that rely on riparian habitat, highlighting the importance of conservation and restoration of these habitats.

### Implications for conservation management

In the savanna landscapes of northern Australia, historical increases in the frequency of high-intensity fires - primarily associated with the breakdown of Indigenous fire management practices following European colonisation - have driven the decline of a range of animals (Woinarski and Legge 2013) and vegetation communities (Russell-Smith et al. 2002); however, the impacts of fire on northern Australian riparian habitats, and the biodiversity they host, have received too little attention. While riparian habitat can provide refuge from wildfires in the surrounding savanna matrix, it can also become a corridor for fire under certain conditions (Pettit and Naiman 2007). Moreover, extended droughts increase the risk of high-severity riparian fires, as do large flood events, which result in the accumulation of woody debris (i.e. fuel) in riparian zones (Pettit and Naiman 2005, 2007). Once a wildfire has occurred in the riparian zone, the subsequent build-up of dead wood also increases the risk of another fire (Pettit and Naiman 2007). Under continued climate change, rainfall is predicted to become more erratic across much of Australia, with more frequent periods of drought as well as more extreme rainfall events leading to large floods (Bureau of Meteorology and CSIRO 2020, Almazroui et al. 2021); hence, unless action is taken, we will see more frequent and severe fire in riparian zones.

Although future research should validate optimal fire frequency and strategy for riparian zones, we can use our current understanding of fire combined with the findings from this study to make recommendations to improve protection of riparian habitat from fire. With the increasing threat of fire to riparian zones, it appears that attempting to exclude fire from riparian habitat altogether constitutes a risky strategy, as the associated accumulation of fuel can lead to an increased risk of| wildfires (Pettit and Naiman 2005, 2007). While reducing fuel loads in savannas adjacent to riparian zones may reduce the likelihood of wildfire impacting riparian zones, later in the dry season, when moisture levels are lower, riparian zones may still become a corridor for fire to travel through (Pettit and Naiman 2007). Hence, the challenge for land managers and conservationists is to minimise the impact of fire when it does enter the riparian zone. We propose that this can be achieved on a landscape scale by using small, low-intensity prescribed burns perpendicularly across the riparian zone to create breaks along riparian corridors, mitigating the risk of large sections of riparian corridors burning in the late dry season. It is however important to implement such burns when conditions are right to prevent escalation to high-intensity fire (i.e. with optimal wind, relatively high moisture levels), and in years when rainfall has been at or above average.

Additionally, fine-scale fire management may be implemented at priority sites to offer maximal protection for high-quality riparian habitat with high purple-crowned fairy-wren density. Here, savanna habitat parallel to the riparian zone may be burnt when savanna habitat has cured but riparian habitat has not yet, followed later by burning small sections of riparian habitat across the waterway once conditions are optimal, thereby creating a complete protective buffer around the high-quality site. Based on the results of this study, this strategy of implementing low-intensity prescribed burns is not expected to affect survival of adult fairy-wrens, which appears important for post-fire population recovery: recovery after a low-intensity fire, with no mortality, was faster, despite reduced breeding success, than after a high-intensity fire. It is important however to consider the timing of fire when planning prescribed riparian burns: by burning immediately after the main breeding season (around April), the effect on reproductive output may be minimised, impacting breeding success for one season only, the subsequent year post-fire.

## AUTHOR CONTRIBUTIONS

Niki Teunissen and Anne Peters conceived the study; Niki Teunissen collected and analysed data; Niki Teunissen led the writing of the manuscript, with input from Hamish McAlpine, Skye Cameron, Brett Murphy, and Anne Peters. All authors gave final approval for publication.

## ACKNOWLEDGEMENTS

We acknowledge the traditional custodians of the country on which this research was conducted. We thank Michael Roast, Marie Fan, Nataly Hidalgo Aranzamendi, and all volunteers who helped collect data. We thank Australian Wildlife Conservancy and staff at Mornington Wildlife Sanctuary for their contributions to fire management and ongoing support. This study was supported by the Australian Research Council (FT110100505; DP150103595, DP180100058 to A.P.) and Monash University. Research was approved by the Monash University Animal Ethics Committee; the Western Australian Department of Biodiversity, Conservation and Attractions; the Australian Bird and Bat Banding Scheme; and the Australian Wildlife Conservancy.

## DATA AVAILABILITY STATEMENT

Data will be archived in the Dryad Digital Repository.

