## Supplementary material: Figure S1, Table S1, Table S2, Table S3, Table S4 for "Demographic impacts of low- and high-intensity fire in a riparian savanna bird: implications for ecological fire management"

**SUPPLEMENTARY INFORMATION**

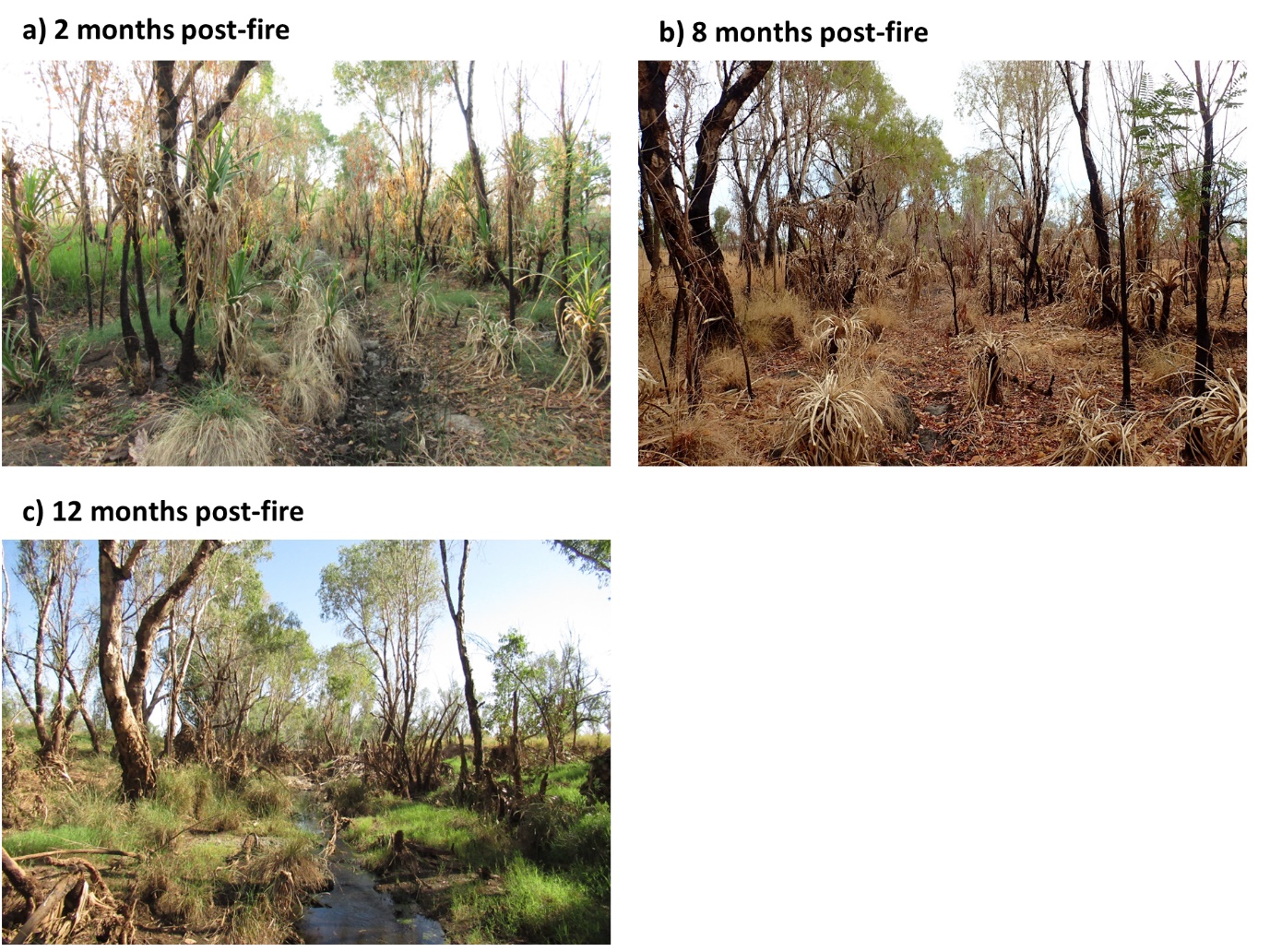

**Fig. S1.** Immediate and delayed effects of intense fire on riparian habitat. Photos of riparian habitat taken at the same location (a) 2 months, (b) 8 months, and (c) 12 months after the intense fire in 2019.

**Table S1.** Full output of the linear models testing for the effect of a low-intensity fire and a high-intensity fire on purple-crowned fairy-wren density (measured as individuals per km of creekline).

|  | **Low-intensity fire** | | | **High-intensity fire** | | |
| --- | --- | --- | --- | --- | --- | --- |
| **Parameter** | ***Β* ± SE** | **t_16_** | ***P*** | ***Β* ± SE** | **t_16_** | ***P*** |
| Intercept | 32.56 ± 2.60 | 12.55 | <0.01 | 17.78 ± 3.05 | 5.83 | <0.01 |
| Time period^a^ | 2.56 ± 3.67 | 0.70 | 0.49 | 19.52 ± 4.31 | 4.53 | <0.01 |
| Fire treatment^b^ | 10.95 ± 3.67 | 2.98 | <0.01 | 11.53 ± 4.31 | 2.67 | 0.02 |
| Time period^a^ x Fire treatment^b^ | -12.04 ± 5.19 | -2.32 | 0.03 | -0.22 ± 6.10 | -0.04 | 0.97 |

^a^ Pre-fire relative to post-fire

^b^ Unburnt relative to burnt

**Table S2.** Full output of the cox proportional hazards models testing for the effect of a low-intensity fire and a high-intensity fire on purple-crowned fairy-wren survival.

|  | **Low-intensity fire** | | | **High-intensity fire** | | |
| --- | --- | --- | --- | --- | --- | --- |
| **Parameter** | ***Β* ± SE** | **z** | ***P*** | ***Β* ± SE** | **z** | ***P*** |
| Fire treatment^a^ | -0.29 ± 0.27 | -1.08 | 0.28 | 0.58 ± 0.20 | -2.75 | <0.01 |
| Status^b^ | 0.12 ± 0.24 | 0.50 | 0.62 | 0.19 ± 0.19 | 1.03 | 0.31 |

^a^ Unburnt relative to burnt

^b^ Subordinate relative to dominant

**Table S3.** Full output of the cox proportional hazards models testing for the effect of a low-intensity fire and a high-intensity fire on purple-crowned fairy-wren’s likelihood to remain at home instead of disperse.

|  | **Low-intensity fire** | | | **High-intensity fire** | | | |
| --- | --- | --- | --- | --- | --- | --- | --- |
| **Parameter** | ***Β* ± SE** | **z** | ***P*** | | ***Β* ± SE** | **z** | ***P*** |
| Fire treatment^a^ | -0.40 ± 0.35 | -1.12 | 0.26 | | 0.22 ± 0.35 | 0.64 | 0.52 |
| Status^b^ | 1.48 ± 0.34 | 4.30 | <0.01 | | 1.60 ± 0.33 | 4.90 | <0.01 |

^a^ Unburnt relative to burnt

^b^ Subordinate relative to dominant

**Table S4.** Full output of the generalised linear mixed models testing for the effect of a low-intensity fire and a high-intensity fire on purple-crowned fairy-wren breeding success.

|  | **Low-intensity fire** | | | **High-intensity fire** | | |
| --- | --- | --- | --- | --- | --- | --- |
| **Parameter** | ***Β* ± SE** | **z** | ***P*** | ***Β* ± SE** | **z** | ***P*** |
| Intercept | -0.46 ± 0.28 | -1.66 | 0.10 | -0.54 ± 0.27 | -2.02 | 0.04 |
| Time period^a^ | 0.77 ± 0.32 | 2.42 | 0.02 | 0.47 ± 0.32 | 1.47 | 0.14 |
| Fire treatment^b^ | 0.49 ± 0.30 | 1.64 | 0.10 | -0.12 ± 0.30 | -0.40 | 0.69 |
| Time period^a^ x Fire treatment^b^ | -0.94 ± 0.35 | -2.69 | <0.01 | 0.11 ± 0.37 | 0.29 | 0.77 |

^a^ Pre-fire relative to post-fire

^b^ Unburnt relative to burnt
